# Outbreak of Western equine encephalitis virus infection associated with neurological disease in horses following a nearly 40-year intermission period in Argentina

**DOI:** 10.1101/2024.09.06.611705

**Authors:** Maria Aldana Vissani, Florencia Alamos, Maria Silvia Tordoya, Leonardo Minatel, Juan Manuel Schammas, María José Dus Santos, Karina Trono, Maria E. Barrandeguy, Udeni B.R. Balasuriya, Mariano Carossino

## Abstract

Western equine encephalitis virus (WEEV) is a mosquito-borne arbovirus (genus *Alphavirus*, family *Togaviridae*) that has reemerged in South America in late 2023 causing severe disease in both horses and humans after a nearly 40-year intermission period. We describe the virological, serological, pathological and molecular features of WEEV infection in horses during the 2023- 2024 outbreak in Argentina. WEEV-infected horses developed neurological signs with mild to severe encephalitis associated with minimal to abundant WEEV-infected cells as demonstrated by WEEV-specific *in situ* hybridization. The distribution of viral RNA was multifocal with predominance within neuronal bodies, neuronal processes, and glial cells in the medulla oblongata and thalamic regions. Phylogenetic analysis of partial nsP4 sequences from three viral isolates obtained from three different provinces of Argentina support grouping with other temporally current WEEV strains from Uruguay and Brazil under a recently proposed novel lineage.

## INTRODUCTION

Western equine encephalitis virus (WEEV) is a mosquito-borne arbovirus and an important cause of severe and often fatal neurologic disease (encephalomyelitis) in both horses and humans in the Americas [1–4]. WEEV belongs to the genus *Alphavirus*, family *Togaviridae*, which also includes the related mosquito-borne Eastern equine encephalitis (EEEV) and Venezuelan equine encephalitis viruses (VEEV) [5]. These are small, enveloped viruses with a positive-sense, single-stranded RNA genome of approximately 11.5kb in length that includes two (non-structural and structural) open reading frames (ORF) flanked by untranslated regions at the 5’ and 3’ ends [5–7]. The non-structural ORF encodes four nonstructural proteins (NSP1- 4), while the structural ORF encodes the structural proteins CP, E3, E2, 6K, TF and E1 [5–7]. The glycoproteins E1 and E2 are expressed on the viral envelope, are strongly immunogenic and induce the production of neutralizing antibodies [8]. Genomic analysis has demonstrated that WEEV is a descendant of an ancient recombination event between Sindbis virus (SINV)- like and EEEV-like ancestors [9, 10], and recent phylogenetic analysis of full-genome sequences of WEEV proposed its classification into two lineages (group A and B), with three sublineages among group B viruses (sublineages B1-B3) [11].

WEEV circulates enzootically in the Americas in a cycle between passerine birds and ornithophilic mosquitoes, being *Culex tarsalis* its primary mosquito vector in North America and is associated with irrigated agriculture in the western US [12]. Other bridging mosquito species can facilitate transmission (e.g. *Ochlerotatus melanimon*, *Aedes dorsalis* and *Aedes campestris*) [12]. In Argentina, the genus *Mansonia* sp has been associated with the epizootic of WEE during 1982-1983 as well as *Anopheles albitarsis* and *Psorophora pallescens* [13], and *Aedes albifasciatus* and *Culex pipiens* complex mosquitoes were demonstrated to be susceptible to WEEV [14, 15]. Small mammals can also participate in a secondary cycle [16] and, interestingly, most Argentinean mosquitoes from which WEEV has been isolated feed principally on mammals, including rice rats (*Oryzomys* sp.) [17] and European hares [13]. Spillover transmission to humans and horses occurs when the enzootic transmission cycle is disrupted, causing outbreaks of disease; both humans and horses serve as dead-end hosts [18]. The disease caused by EEEV, WEEV, and VEEV is clinically indistinguishable, and is characterized by fever, anorexia, depression and clinical signs of encephalomyelitis, and case fatality is typically high [3, 19–21].

WEEV was first isolated in 1930 in California, U.S. from the brain of an encephalitic horse [22]. During the 1930’s through the 1950’s, WEEV produced widespread outbreaks in the western U.S. and Canada [18]. WEEV infections in North America have drastically decreased, with no human cases registered since 1999 [23]. Furthermore, the virus has not been detected in mosquito pools since 2008 [11].

In Argentina, WEEV was isolated for the first time from horses with neurologic disease in 1933. The major WEEV outbreaks in horses occurred in 1972-1973 and 1982-1983 [24–26], with two and five human cases registered in 1973 and 1983, respectively [27]. Additionally, serologically positive horses were detected via surveillance between 1983 and 1985 [24]. The last case of WEE (human) in Argentina was detected in 1996 [28], and the last one in the region occurred in 2009 in Uruguay [29]. Vaccines commercially available in Argentina against EEEV and WEEV for use in horses include two bivalent inactivated vaccines containing EEEV and WEEV (Tecnovax and Rosenbusch, Buenos Aires, Argentina) and two polyvalent inactivated vaccine containing WEEV/EEEV (Tecnovax and Zoetis Animal Health, Kalamazoo, MI, USA). While mandatory until 2016, since then vaccination against EEEV and WEEV became non-mandatory per regulations from the national animal health authorities (SENASA Resolution No. 521/16).

During late November 2023, an epizootic of WEE in horses was reported in Argentina after a nearly 40-year intermission period following the 1982-1983 outbreak [30]. To date, a total of 1,530 cases (including suspected and confirmed cases) have been recorded in 17 provinces. Within the framework of the WEEV health emergency established by Resolution 1219 /2023, mandatory vaccination against EEEV/WEEV in Argentina was re-established in January 2024 for all horses that are at least 2-month-old. Here, we report the virological, serological, pathological, and phylogenetic features of neurologic cases received by the Equine Virology Unit, INTA during the 2023-2024 WEEV epizootic in the horse population from Argentina.

## MATERIALS AND METHODS

### Samples and processing

Samples from a total of 34 horses showing neurologic disease were submitted to the Equine Virology Unit, Institute of Virology, CICVyA, INTA, Hurlingham, Argentina. All the clinical cases from which samples were received had been communicated to the national animal health authorities (SENASA). Samples submitted from these cases included brain (n=32), other visceral organs (n=33) and cerebrospinal fluid (CSF; n=19). Tissue samples fixed in 10% neutral buffered formalin (brain, lung, liver and/or spleen) from six of these horses and serum samples from 49 unvaccinated horses related to the outbreak were also submitted. Upon receipt, samples were processed in a biosafety level 4 laboratory (BSL-4) and 10% tissue homogenates were prepared by homogenizing 1g of tissue in 9 ml of complete DMEM using a mortar and pestle in a biosafety cabinet (class II). Tissue homogenates were clarified by centrifugation at 2,500 X g for 10 minutes at 4 °C; the resultant supernatants were subsequently collected and stored at -80 °C until used. Serum samples were aliquoted and stored at -20 °C until analyzed. Additionally, tissue samples (brain) from six non-equid species (ovine [n=3], porcine [n=1], and cervid [n=2]) showing neurologic signs were also submitted.

### Cells and media

Vero E6 cells (CRL-1586, American Tissue Type Collection [ATCC], Manassas, VA, USA) were maintained in Dulbecco’s Modified Minimum Essential Medium (DMEM) containing 10% heat-inactivated fetal bovine serum, 200 mM L-glutamine, penicillin and streptomycin (100 U/ml and 100 μg/ml) and 0.25 μg/ml of amphotericin B (INTA, Hurlingham, Argentina). Vero E6 cells up to passage 50 were used for virus isolation and virus neutralization tests.

### Nucleic acid isolation

Viral RNA was extracted from 10% tissue homogenates and CSF using the QIAamp Viral RNA Mini Kit (QIAGEN, Cat. No. 52904, Valencia, CA, USA), in accordance with the manufacturer’s instructions and stored at -80 °C until used.

### Pan-Alphavirus genus reverse-transcription nested polymerase chain reaction

For initial diagnosis, a pan-Alphavirus genus reverse-transcription, nested polymerase chain reaction (RT-nPCR) targeting a 481 bp region of the nsP4 gene was performed using previously described primer sets [31]. The primers 1+ and 1-, and 2+ and 2- for the first and second PCR rounds were used, respectively, as previously described. Extracted nucleic acids were first subjected to a reverse transcription (RT) reaction using the High-Capacity cDNA Reverse Transcription kit (ThermoFisher Scientific, Cat N° 4368813, Vilnius, Lithuania). Briefly, the RT reaction mix was composed of 3 μl of 10X RT buffer, 1.2 μl of 25X dNTP mix, 3 μl of 10X RT random primers, 1.5 μl of the MultiScribe Reverse Transcriptase (50 U/μl), 11.3 μl of nuclease-free water and 10 μl of extracted nucleic acid. The reaction included 10 min at 25 °C and 45 min at 37 °C. Following the RT step, the cDNA was subjected to the first round of PCR. The 25 μl reaction contained 5 μl of PCR Green Buffer with magnesium chloride (Promega, Cat N° M7845, Madison, USA), 1 μl of 5 mM dNTP, 0.75 μl of 50 mM magnesium chloride, 0.25 μl of GoTaq DNA polymerase 2.5 U/ml (Promega, Cat N° M7845, Madison, USA), 11 μl of nuclease-free water, 0.4 μM of each primer (1+ and 1-), and 5 μl of cDNA. The cycling parameters for the first PCR round included: 5 min at 94 °C, 40 cycles at 94 °C for 30 sec, 52 °C for 30 sec, and 72 °C for 30 sec; and a final extension at 72 °C for 7 min. The initial PCR product (481 bp) was subjected to a second round of PCR with a 25 μl reaction similar to the first PCR round except for 0.25 μl of 50 mM magnesium chloride, 15.5 μl of nuclease-free water, 0.4 μM of primers 2+ and 2-, and 1 μl of the PCR product as template. The cycling conditions were equivalent to the first round of PCR. RNA extracted from a chimeric Sindbis strain kindly provided by Dr. Contigiani (Vanella Institute, Córdoba University, Cordoba, Argentina) was used as a positive control. The final PCR products (195 bp) were electrophoretically separated on a 2% agarose gel supplemented with Sybr™ Safe DNA Gel Stain (Invitrogen by ThermoFisher Scientific, Cat S33102, Carlsbad, California, USA) and visualized under UV light. All samples submitted were also subjected to in-house equine herpesvirus type 1 (EHV-1) and West Nile virus (WNV)-specific standard PCR and RT-PCR, respectively, to rule out these possible differential diagnoses.

### EEEV/WEEV-specific reverse-transcription TaqMan^®^ real-time polymerase chain reaction

On samples that resulted positive to the pan-Alphavirus genus RT-nPCR, nucleic acids were subjected to a EEEV or WEEV-specific TaqMan^®^ RT-qPCR targeting the E2 and E1 gene, respectively, using the primer and probe sets as previously described [32] and the AgPath-ID™ One-Step RT-PCR mix (ThermoFisher Scientific, Cat N4387424, USA). Briefly, 25 μl reaction containing 12.5 μl of 2X AgPath-ID™ One-Step RT-PCR mix, 1 μl 25X RT Mix, 0.2 μM of the EEEV or WEEV-specific TaqMan^®^ fluorogenic probe (FAM-TAMRA), 0.4 μM of each primer, 5.8 μl of nuclease-free water, and 5 μl of template RNA. An ABI 7500 Fast Real-time PCR System (Applied Biosystems^®^) was used with the following program: 10 min at 45 °C (reverse transcription step), 10 min at 95 °C (PCR initial activation step), and 45 cycles at 95 °C for 15 sec (denaturation) and 60 °C for 60 sec (combined annealing/extension). A control plasmid containing the respective target sequences kindly provided by Agence Nationale Sécurité Sanitaire Alimentaire Nationale (ANSES), Maisons-Alfort, France was used as positive control. The cutoff value was determined to be a cycle threshold value of 40.

### Virus isolation

Virus isolation was attempted in samples confirmed positive to WEEV via RT-qPCR and performed within a BSL-4 laboratory. One ml of 10% brain homogenates were inoculated onto confluent Vero E6 monolayers grown in 25-cm^2^ flasks and adsorbed for 1 hour at 37°C. Following adsorption, monolayers were washed twice with complete DMEM, and 9 ml of fresh complete DMEM added. Inoculated flasks were incubated at 37°C, 5% CO2 for 7 days and reviewed for signs of cytopathic effect (CPE) daily for 7 days followed by 3 blind passages if no CPE was observed. Positive results were determined by the presence of CPE and were confirmed using the WEEV-specific real time RT-qPCR assay from nucleic acids extracted from tissue culture supernatants. Virus isolates were titrated in Vero E6 cells, titers calculated using the Reed and Muench method and expressed as tissue culture infectious dose 50% per ml (TCID50/ml) and stored at -80 °C.

### Sequencing of nsP4 and phylogenetic analysis

For Sanger sequencing, a 481 bp region of the nsP4 gene was amplified as indicated above from nucleic acids derived from three virus isolates obtained in this study (E5191.23.3P, E5239.23.1P, and E5288.23.1P, respectively). The primers 1+ and 1- were used for RT-PCR and Sanger sequencing at the Instituto de Biotecnología sequencing service, Instituto Nacional de Tecnología Agropecuaria (INTA, Buenos Aires, Argentina). Sequences obtained were analyzed using Geneious Prime (Dotmatics, Boston, MA, USA) and subjected to multiple sequence alignment with other WEEV sequences available in GenBank using MUSCLE [33] on MEGA 11. Genetic distances were calculated using the Kimura 2-parameter+I (invariant sites) model at the nucleotide level, and the phylogenetic trees were constructed using the Maximum-likelihood method with 1,000 bootstrap replicates on MEGA11 [34]. The nsP4 gene of the RefSeq for EEEV (North American variant, GenBank accession NC_003899) was used as an outgroup. The phylogenetic tree was visualized and edited using TreeViewer [35]. Sequences obtained in the current study were deposited in GenBank under accession numbers: 1009, strain E.5191.23.3P; 1010, strain E.5239.23.1P; and 1011, strain E.5288.23.1P.

#### Virus neutralization test

Serum samples were analyzed for WEEV-specific neutralizing antibodies using a virus (micro)neutralization test (VNT) performed in a BSL-4 laboratory. A total of 49 serum samples were tested, these included (A) horses that succumb to neurologic disease and were confirmed as WEEV-infected via real time RT-qPCR (n=8), (B) horses with neurologic disease that cohabitated with a horse that previously died and was confirmed as WEEV-infected via real time RT-qPCR (n=9), (C) horses without neurologic disease that cohabitated with a horse that previously died and was confirmed as WEEV-infected via real time RT-qPCR (n=6), (D) horses with neurologic signs but with no WEEV-confirmed cases in the premises (n=23), and (E) convalescent horses that developed neurologic disease and recovered (n=3). Among this sample set, there were also 4 horses within group D that were serologically monitored for up to 3 months following initiation of neurologic signs. As mentioned, all serum samples were obtained from horses with no history of previous vaccination. From samples of CSF, 18 were also subjected to VNT, 8 of which had their corresponding serum samples. Briefly, serial two-fold dilutions (1:5 to 1:640) of heat-inactivated serum or CSF samples were prepared in triplicate in 50 μl of complete DMEM. Fifty microliters of a working WEEV suspension (E.5191.23 Q/P 4P 90124) containing 100 tissue culture infective doses 50% per 50 μl (100 TCID50/50 μl) was added to each well, and the plates were incubated for 1.5 h at 37 °C in 5% CO2. Finally, 100 μl/well of a suspension of Vero E6 cells at a concentration of 2x10^5^ cells/ml were added, and the plates were incubated at 37 °C in 5% CO2 for 72 h. The neutralizing antibody titer was recorded as the reciprocal of the highest serum dilution that provided at least 50% neutralization of the cytopathic effect produced by the reference virus.

#### Histopathology

The encephalon of 6 horses was dissected and fixed in 10% neutral buffered formalin for up to 14 days. Sections from the cerebral cortex (frontal, parietal and occipital lobes), thalamus, internal capsule, mesencephalon, cerebellum, pons, medulla oblongata, and cervical spinal cord were obtained from each case, processed routinely, and embedded in paraffin. Subsequently, four-micron tissue sections were obtained and stained with hematoxylin and eosin following standard histological procedures.

### WEEV-specific RNAscope^®^ *in situ* hybridization (ISH)

For WEEV-specific RNAscope^®^ ISH, an anti-sense probe targeting the nucleotides 147 – 1964 of the polyprotein gene of the WEEV McMillan strain genome (GenBank accession number AF229608.1) was designed and synthesized by Advanced Cell Diagnostics (Catalog number 493418, ACD, Newark, CA). The RNAscope® ISH assay was performed using the RNAscope 2.5 LSx Reagent Kit (Advanced Cell Diagnostics, Newark, CA) on the automated BOND RXm platform (Leica Biosystems, Buffalo Grove, IL) as described previously [36]. Briefly, four-micron sections of formalin-fixed paraffin-embedded tissues were mounted on positively charged Superfrost^®^ Plus Slides (VWR, Radnor, PA) and subjected to automated baking and deparaffinization followed by heat-induced epitope retrieval (HIER) using a ready-to-use EDTA-based solution (pH 9.0; Leica Biosystems) at 100 °C for 15 min. Subsequently, tissue sections were treated with a ready-to-use protease (RNAscope^®^ 2.5 LSx Protease) for 15 min at 40 °C followed by a ready-to-use hydrogen peroxide solution for 10 min at room temperature. Slides were then incubated with the ready-to-use probe mixture for 2 h at 40 °C, and the signal amplified using a specific set of amplifiers (AMP1 through AMP6 as recommended by the manufacturer). The signal was detected using a Fast Red solution for 10 minutes at room temperature. Finally, slides were counterstained with a ready-to-use hematoxylin for 5 min, followed by five washes with 1X BOND Wash Solution (Leica Biosystems). Slides were rinsed in deionized water, dried in a 60 °C oven for 30 min, and mounted with Ecomount® (Biocare, Concord, CA, USA). As a negative control, sections of FFPE brainstem from an EEEV-infected horse were used.

#### Whole slide scanning and quantitative image analysis

RNAscope^®^ ISH slides were scanned at 40X magnification using an Akoya PhenoImager HT whole slide scanner (Akoya Biosciences, Cambridge, MA, USA). Quantitative analysis was performed in QuPath 0.5.1 digital pathology image analysis software [37]. Briefly, stain vectors were adjusted for each slide respectively before pursuing further analysis. Subsequently, a thresholder for pixel classification was created to automatically detect tissue areas on the slide for analysis. This was achieved following QuPath tutorial guides. Following automated tissue detection, positive pixel counting was performed by setting a positive threshold for the FastRed chromogen following QuPath tutorial guides. The positively stained area was expressed as a percentage of total tissue area quantified and exported into an Excel file.

#### Statistical analysis

Data distribution was evaluated using JMP17 Pro (Cary, NC). Non-parametric correlation analysis between Ct values and percentage ISH positive area was performed using the Spearman’s test on JMP17 Pro. Graphics were subsequently generated using either JMP17 Pro or GraphPad Prism 9 (GraphPad Software, San Diego, CA). The level of significance was set at *P*<0.05 for all tests.

## RESULTS

### Diagnosis of WEEV and virus isolation

WEEV RNA was detected via RT-nPCR and real time RT-qPCR in a total of 25 out of 34 equine case submissions (74%). Specifically, WEEV RNA was detected in brain tissue (n=33) and in the lung from a single case, with Ct values ranging from 26.5 to 38.2. WEEV RNA was not detected in any of the CSF samples received (n=19) even though WEEV RNA was detected in the brain in all of those instances. WEEV-positive samples derived from cases that occurred in eight different provinces (Buenos Aires [9], Entre Ríos [4], La Pampa [3], Mendoza [4], Corrientes [2], Río Negro [1], Santa Fé [1] and Chubut [1], Figure 1), without epidemiological relationship between each other. All the horses in which WEEV infection was confirmed were working horses that had no history of previous vaccination. All samples (positive or negative for WEEV) were negative for EHV-1 DNA and WNV RNA. Interestingly, from the non-equid cases received, WEEV was detected in the brain of a sheep derived from Buenos Aires province.

**Figure 1.**
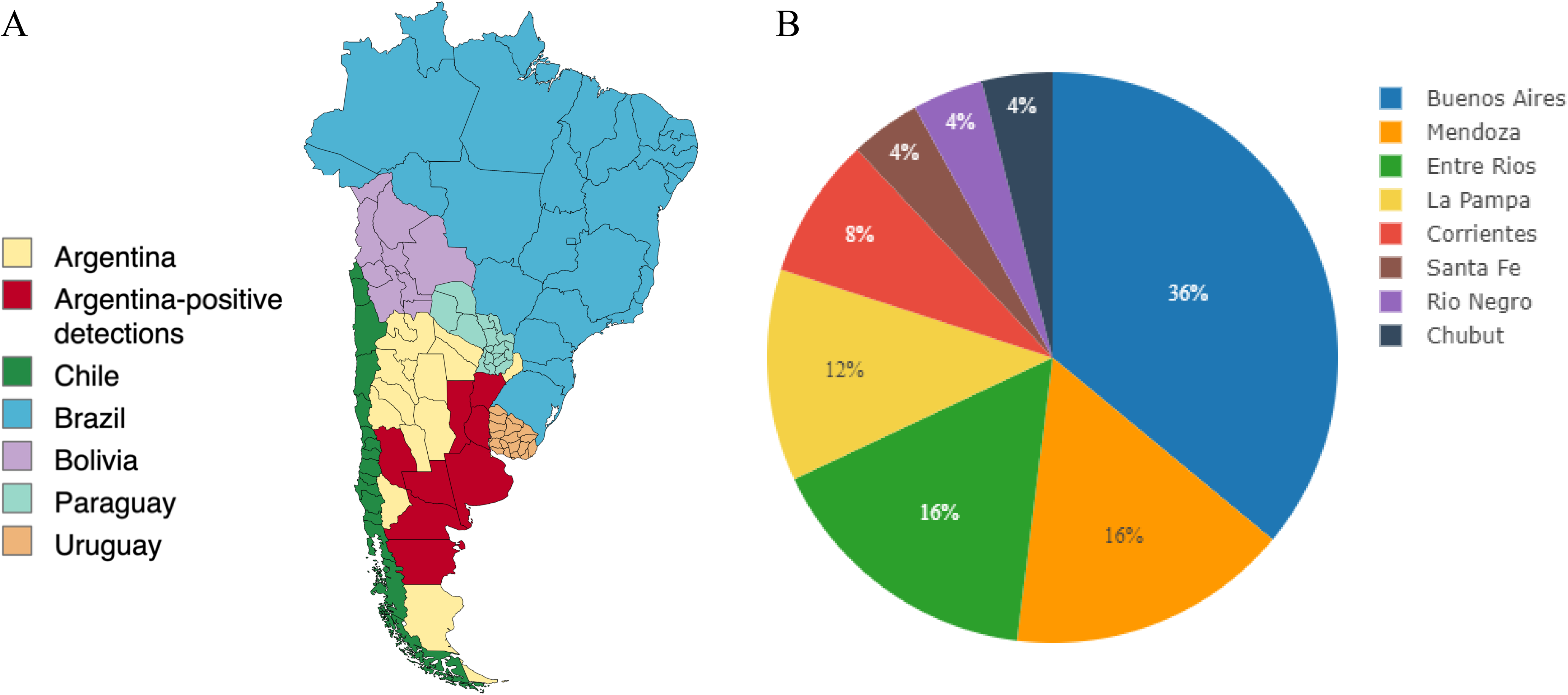
Geographical distribution of WEE cases included in this study. (A) Cases included derived from 8 provinces from Argentina (marked in red). (B) Case distribution by province.

Virus isolation was attempted from seven samples derived from five of the twenty-five WEEV- positive cases based on the availability of sufficient brain homogenate (Table 1). Following inoculation into Vero E6 cells, cytopathic effect (CPE) was noticed in 5/7 inoculated tissue samples derived from 4/5 WEEV-positive cases originating from Buenos Aires, Mendoza, Corrientes and Entre Rios provinces, respectively. Three of the isolates showed evidence of CPE at the first passage while one showed evident CPE only at the third blind passage. Titers for the isolates obtained after a single passage ranged between 10^6.33^ to 10^7^ TCID50/ml, while the titer of the single sample that required three blind passages was 10^6.66^ TCID50/ml. The identities of the four virus isolates were confirmed by WEEV-specific RT-qPCR, with Ct values ranging from 21.25 to 24.7.

**Table 1.**
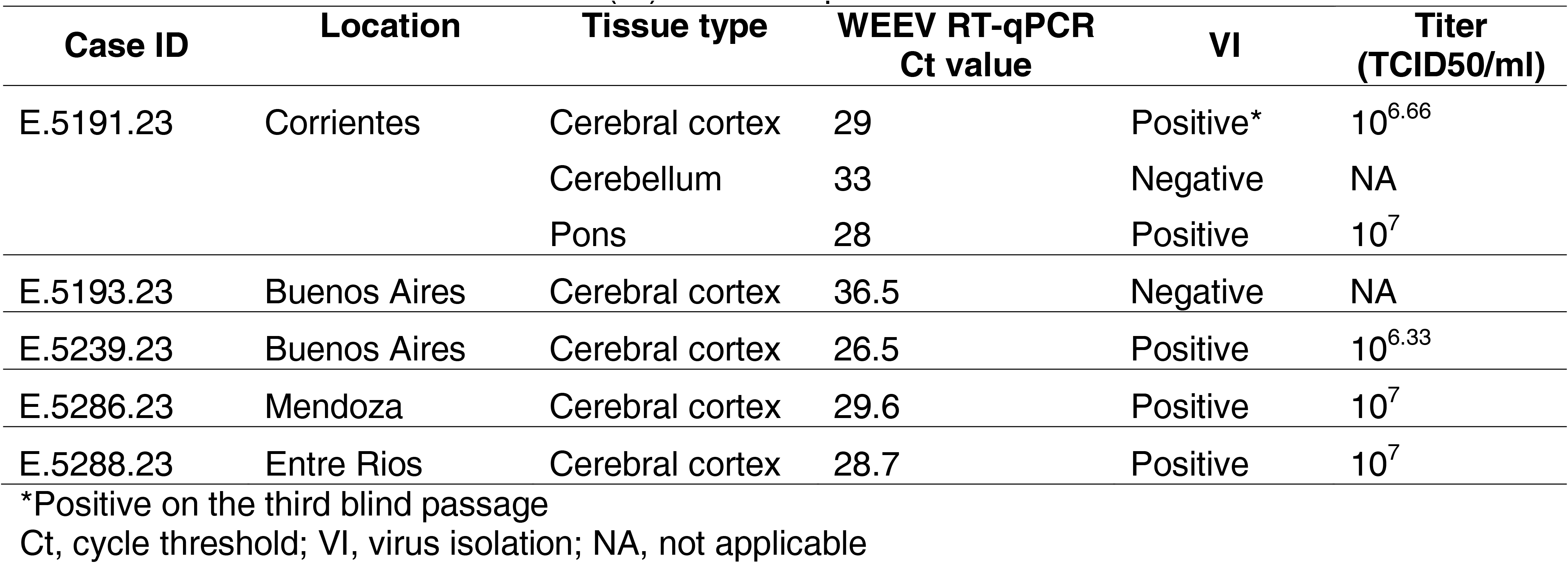
Cases for which virus isolation (VI) was attempted

| Case ID | Location | Tissue type | WEEV RT-qPCR<br>Ct value | VI | Titer<br>(TCID50/ml) |
| --- | --- | --- | --- | --- | --- |
| E.5191.23 | Corrientes | Cerebral cortex | 29 | Positive* | $10^{6.66}$ |
|  |  | Cerebellum | 33 | Negative | NA |
| | | Pons | 28 | Positive | $10^7$ |
| E.5193.23 | Buenos Aires | Cerebral cortex | 36.5 | Negative | NA |
| E.5239.23 | Buenos Aires | Cerebral cortex | 26.5 | Positive | $10^{6.33}$ |
| E.5286.23 | Mendoza | Cerebral cortex | 29.6 | Positive | $10^7$ |
| E.5288.23 | Entre Rios | Cerebral cortex | 28.7 | Positive | $10^7$ |

\*Positive on the third blind passage
Ct, cycle threshold; VI, virus isolation; NA, not applicable

### Phylogenetic analysis of partial nsP4 gene sequences

Partial nsP4 gene sequences derived from 3/5 virus isolates obtained from infected horses were phylogenetically compared with a total of 50 WEEV sequences available in GenBank (Figure 2). Phylogenetic analysis demonstrated that WEEV strains from South America, including the three WEEV isolates obtained in this study, group into a single clade along with other current WEEV strains from 2023 and 2024 from Brazil and Uruguay, likely belonging to the novel WEEV C lineage proposed recently and most closely related to the CBA87 isolate dated from 1958. Sequence reads derived from the two (E.5193.23 and E.5286.23) of the five isolated strains were not of sufficient quality for analysis.

**Figure 2.**
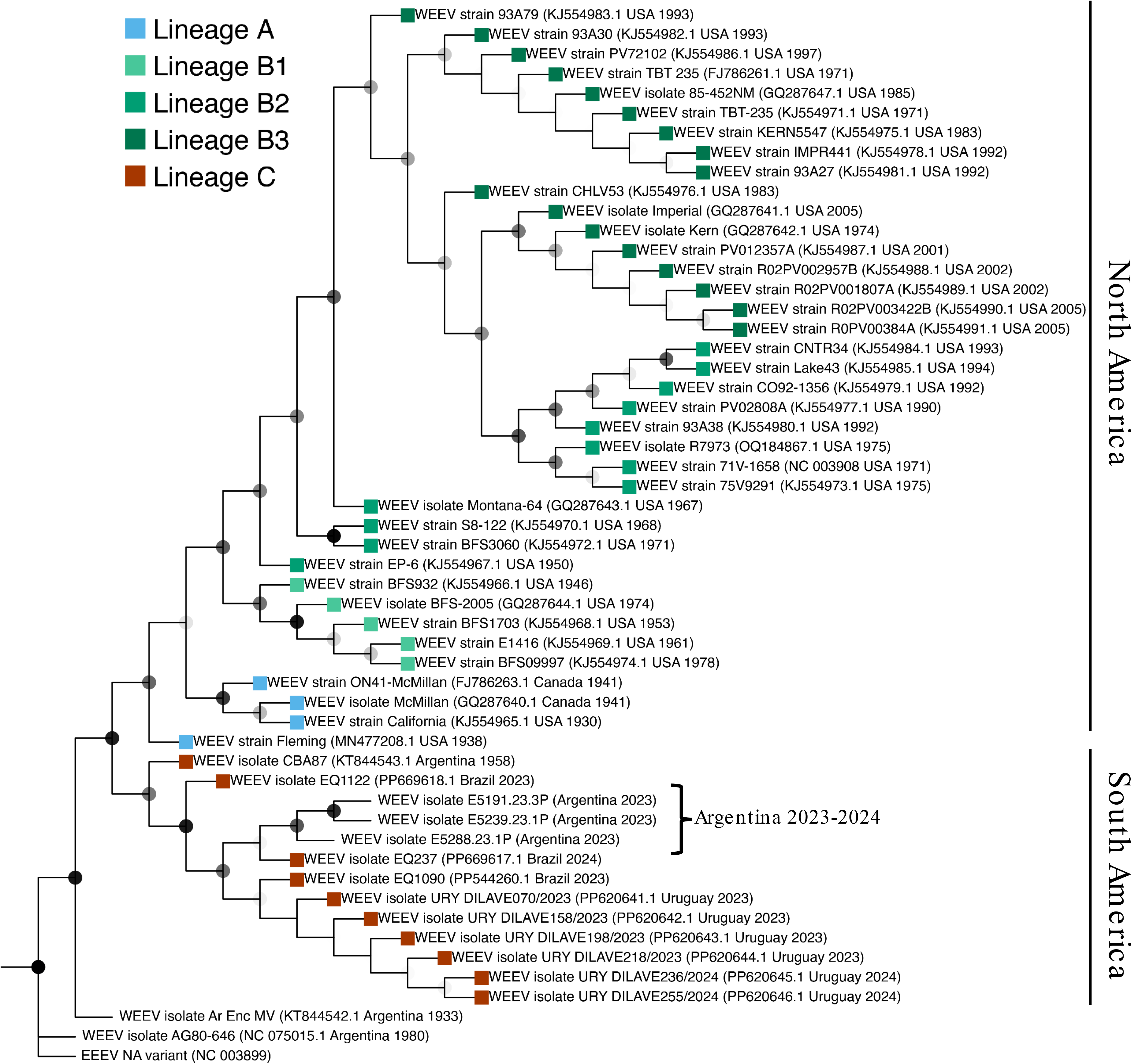
Phylogenetic analysis of partial nsP4 nucleotide sequences of three WEEV isolates from horses from Argentina (marked with a bracket). Current WEEV strains group closely with other reported strains from Uruguay and Brazil (lineage C, red leaves). Maximum likelihood trees were constructed using partial nucleotide sequences and the tree was edited with TreeViewer. A North American variant of EEEV was used as an outgroup. Bootstrap values (>50%) for 1,000 replicates are shown as nodes in grades of gray where black nodes show the highest support (>90%).

### Serological responses to WEEV in infected, comingling, and convalescent horses

Serum samples from 49 horses were analyzed by VNT. Samples were classified into five groups as follows: group A (n=8), horses that died of neurologic disease and had a confirmed postmortem diagnosis of WEEV; group B (n=9), horses with neurological signs that were comingling with a WEEV-confirmed case; group C (n=6), horses without neurological signs that comingled with a WEEV-confirmed case; group D (n=23), horses with neurological signs but no confirmed cases of WEEV in the premises; and group E (n=3), horses that presented with neurological signs in premises with WEEV-diagnosed cases and subsequently recovered (convalescent). A summary of the serological findings per group are shown in Table 2. Importantly, most of the samples under group A (75%) had titers >1:10 and WEEV antibodies were detected in 73.3% of the serum samples derived from comingling horses with and without neurological signs combined (groups B and C). Additionally, we were able to collect three consecutive serum samples from a total of four recovered horses that had clinical evidence of neurologic disease at approximately 1-month interval between the first and second blood collection and at a 2-month interval between the second and third blood collection in order to assess seroconversion. During this period, these horses were not vaccinated. Initial neutralizing antibody titers had a geometric mean titer of 226 (range 1:160 to 1:320), with titers declining gradually (geometric mean titers of 135 [range 40- 320) and 57 [range 20-160] for the second and third set of samples, respectively) (Figure 3).

**Figure 3.**
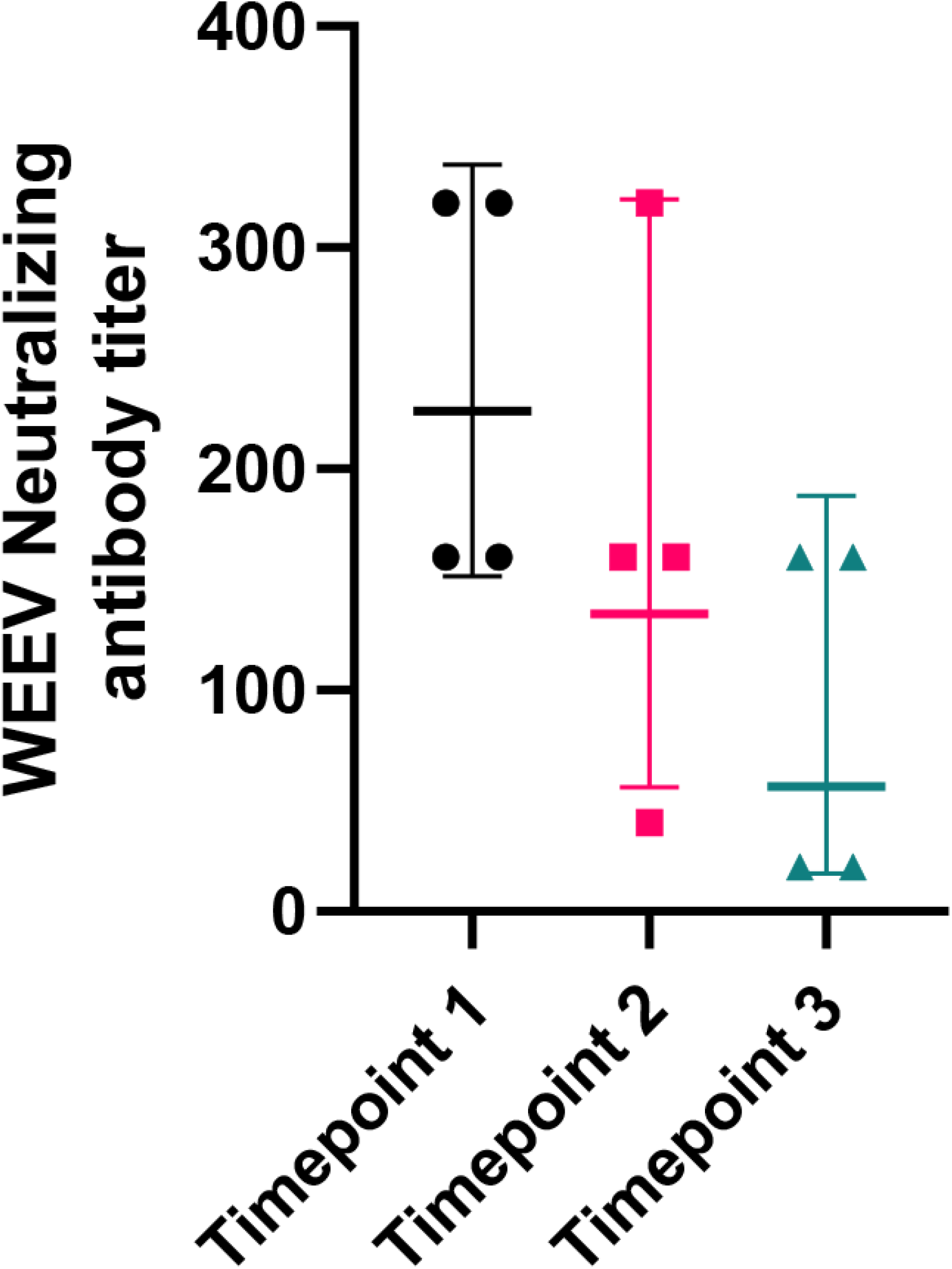
Virus neutralizing antibody dynamics spanning a 3-month period in four horses that recovered from neurologic disease during the 2023-2024 WEEV outbreak. Timepoint 1 and 2 are at a 1-month interval, while timepoints 2 and 3 are at a 2-month interval. Neutralizing antibody titers gradually decline following infection.

**Table 2.** Serological findings in horses during the 2023-2024 WEEV outbreak in Argentina

| **Group** | **n** | **% of samples**<br>**per group** | **% positive**<br>**samples** | **Geometric mean**<br>**titers (range)** |
| --- | --- | --- | --- | --- |
| A: WEEV-infected horses | 8 | 16.3% | 75% (6/8) | 143 (<10-320) |
| B: comingled with neurologic signs | 9 | 18.4% | 100% (9/9) | 173 (80-320) |
| C: comingled without neurologic signs | 6 | 12.2% | 33.3% (2/6) | 28 (<10-40) |
| D: neurologic signs, no WEEV-confirmed<br>cases in premises | 23 | 47.0% | 87% (20/23) | 82 (<10-320) |
| E: convalescent | 3 | 6.1% | 100% (3/3) | 160 (80-320) |

Regarding the CSF samples (n=18), these corresponded to horses classified in groups A (13/18; 72.2%) and D (5/18; 27.8%). Most CSF samples were either negative for anti-WEEV antibodies (11/18; 61.1%) or had a low antibody titer usually between 1:10 to 1:20 (6/18; 33.3%). Among those horses from which both serum and CSF samples were available (8/18), 6 samples had serum neutralizing antibodies and only 50% of these (3/6) had detectable, low titers of antibodies in CSF. Only one CSF sample, corresponding to a horse classified in group A, had a titer of 1:160; however, no paired serum sample was available for this horse.

### Histological lesions, virus tropism and viral abundance in the central nervous system of WEEV-infected horses

The encephalon of a total of 6 WEEV-infected horses (n=5 confirmed via real time RT-qPCR and n=1 confirmed via RNAscope^®^ ISH, see below) was made available for histologic evaluation that included, wherever possible, sections from multiple areas within the encephalon: cerebral cortex, frontal lobe (n=5), cerebral cortex-parietal lobe (n=5), cerebral cortex-occipital lobe (n=5), cerebral cortex-undetermined site (n=1), internal capsule (n=5), thalamus (n=5), mesencephalon (n=4), cerebellum (n=3), pons (n=3), medulla oblongata (n=2) and cervical spinal cord (n=1). In two horses, sections of spleen and liver were also evaluated. In all cases evaluated (n=6), the histologic alterations were consistent with a multifocal to diffuse lymphocytic to mixed meningoencephalitis that was frequently severe (n=4) and occasionally mild (n=1) to moderate (n=1) and affected all examined areas of the encephalon. Histologic alterations affected both the grey and white matter and included perivascular cuffing composed of mostly lymphocytes, and fewer histiocytes and neutrophils infiltrating Virchow-Robin spaces and occasionally extending into the neuroparenchyma, multifocal gliosis, scattered foci of hemorrhage, neuronal degeneration and necrosis featured by shrunken neuronal bodies, chromatolysis, cytoplasmic hypereosinophilia, and nuclear pyknosis; and mild infiltration of the leptomeninges by lymphocytes and histiocytes (Figure 4). In at least one of the cases examined, there were sporadic fibrin thrombi within the vasculature and occasional fibrin exudation into scattered Virchow-Robin spaces, along with intense neutrophilic infiltration of the neuroparenchyma with formation of incipient areas of liquefactive necrosis. While inflammatory lesions were present in all examined areas of the encephalon, they were most frequent and/or intense along the internal capsule, thalamus, and brainstem (pons and medulla oblongata). Similar inflammatory lesions were observed in the cervical spinal cord examined from one of the cases. No significant histologic alterations were noted in the spleen (n=2) and histologic alterations in the liver of two of the cases were characterized by either mild diffuse hepatic lipidosis or mild diffuse periportal fibrosis, both changes considered to be unrelated with WEEV infection.

**Figure 4.**
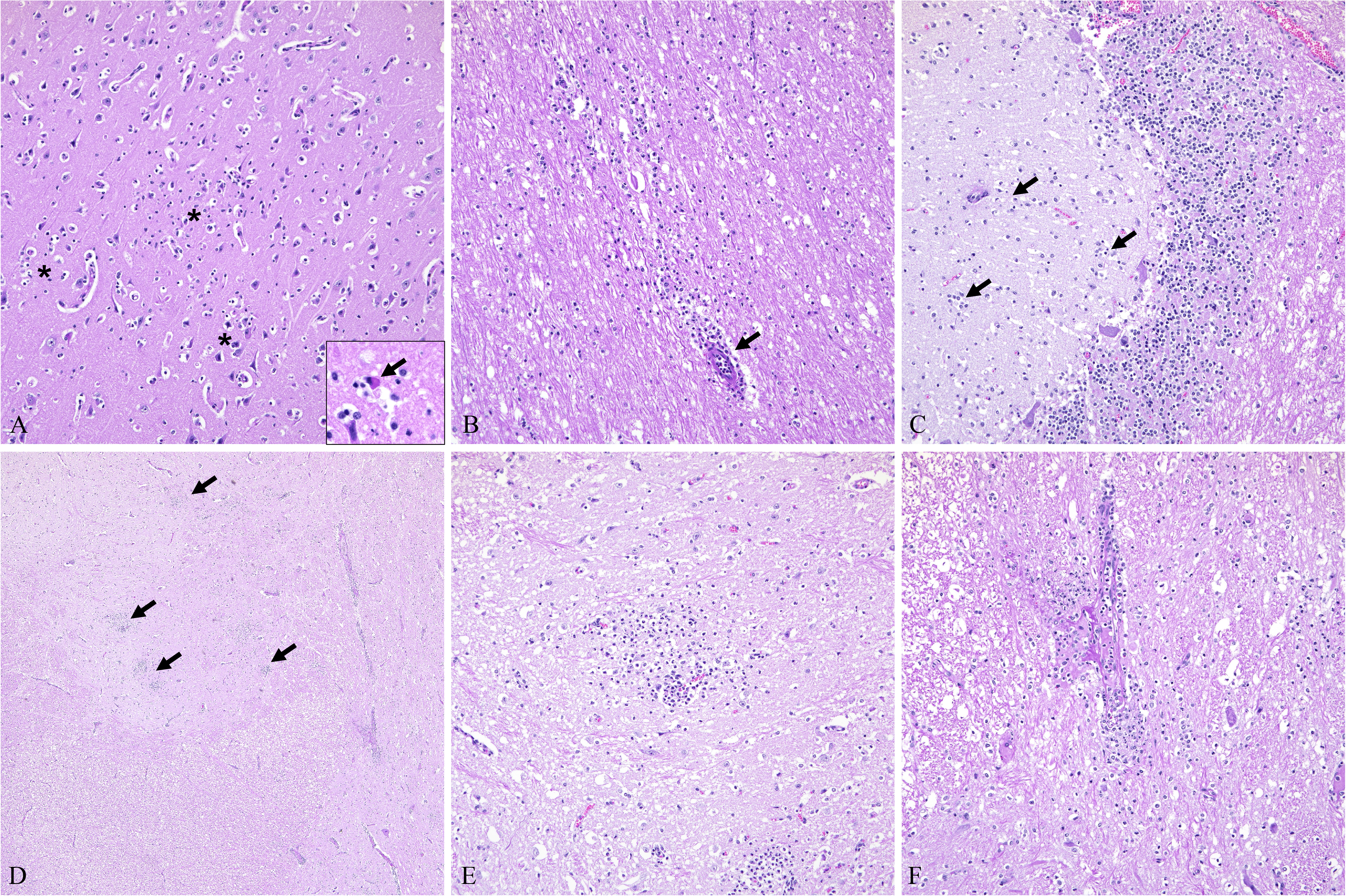
Histological lesions in the CNS associated with WEEV infection in horses. (A and B) In the cerebral cortex, there are multiple foci of gliosis within the grey (A, asterisks) and white matter (B) with occasional neuronal necrosis (A, inset and arrow). Inflammatory cells (predominantly lymphocytes, macrophages and few neutrophils) infiltrate perivascular spaces (B, arrow) as well as the neuroparenchyma. (C) The molecular layer of the cerebellum has multiple foci of gliosis and inflammatory cells (arrows). (D, E and F) The neuroparenchyma of the medulla oblongata has frequent foci of gliosis (arrows) and infiltrating neutrophils (E) with occasional blood vessels affected by fibrinoid change (F). Hematoxylin and eosin. A, B, C, E and F, 200X total magnification; D, 40X total magnification.

RNAscope^®^ ISH using a WEEV-specific probe was used to detect viral RNA *in situ* and determine viral tropism/distribution and viral load. We have analyzed representative sections from the cerebral cortex, thalamus, cerebellum, medulla oblongata, all of which showed variable viral RNA signal in all animals confirmed to be WEEV-infected via RT-qPCR. Additionally, we confirmed WEEV infection in one animal for which only formalin-fixed tissue was available for diagnosis and considered unsuitable for RT-qPCR testing. Viral RNA signal was undetectable in only two examined tissues from a single animal (cerebral cortex and cervical spinal cord). WEEV RNA was frequently detected in areas of white and grey matter and, interestingly, its distribution was multifocal and associated with areas of the neuroparenchyma affected by inflammation and/or neuronal necrosis. Specifically, viral RNA was detected within the cytoplasm of neuronal bodies and their projections, as well as within the cytoplasm of glial cells and infiltrating inflammatory cells in the neuroparenchyma, including the Purkinje cell layer of the cerebellum as well as the cerebellar granular cell layer (Figure 5). No viral RNA was detected in inflammatory cells within Virchow-Robin spaces that formed perivascular cuffs.

**Figure 5.**
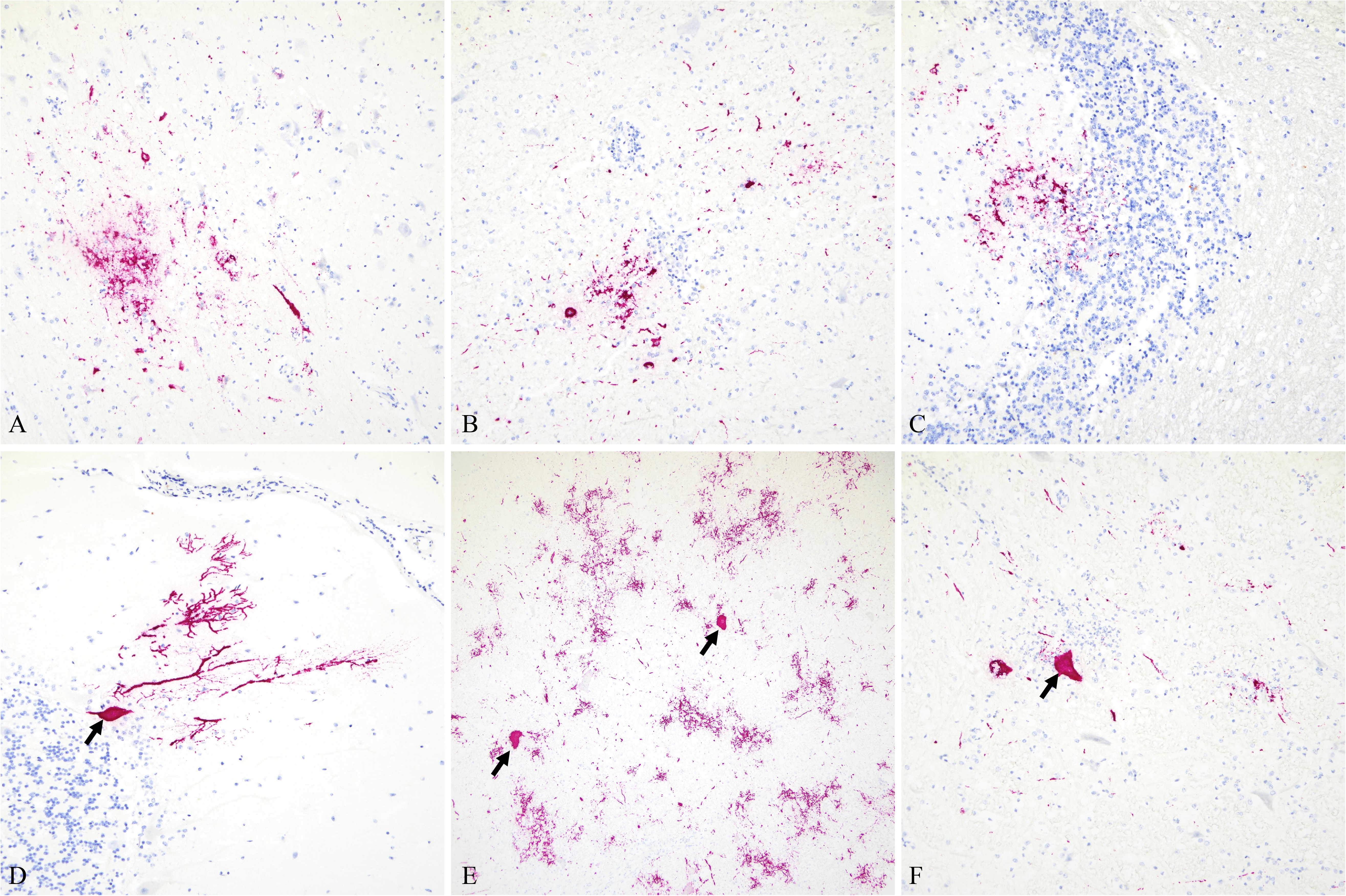
Intralesional detection of WEEV genomic RNA in the CNS of infected horses via RNAscope® *in situ* hybridization. Viral RNA (Fast Red) was detected in all areas of the encephalon examined, including cerebral cortex (A), thalamus (B), cerebellum (C and D), and brainstem (E and F). Overall, viral RNA was detected within the cytoplasm of neurons (D, E and F, arrows) as well as within neuronal projections and glial cells within areas of gliosis. In the cerebellum, viral RNA was detected within the molecular and granular cell layer (C), as well as in Purkinje cells (D). A, B, C, and D, 200X total magnification; E, 40X total magnification; F, 100X total magnification.

Quantitative analysis was performed to correlate WEEV-specific ISH signal with RT-qPCR data, and to specifically map areas of the encephalon based on viral RNA abundance (Table 3 and Figure 6). Both WEEV-specific ISH and Ct values were significantly inversely correlated (Spearmans’ ρ = -0.81, *P* = 0.0149), thus, indicating positive correlation between RNA ISH signal and viral genomic RNA load determined by RT-qPCR. Importantly, WEEV genomic RNA was most abundant in sections of the brainstem followed by the thalamus and was least abundant in the cerebellum and cerebral cortex even though these differences were not statistically significant (*P* = 0.2996; Figure 6).

**Figure 6.**
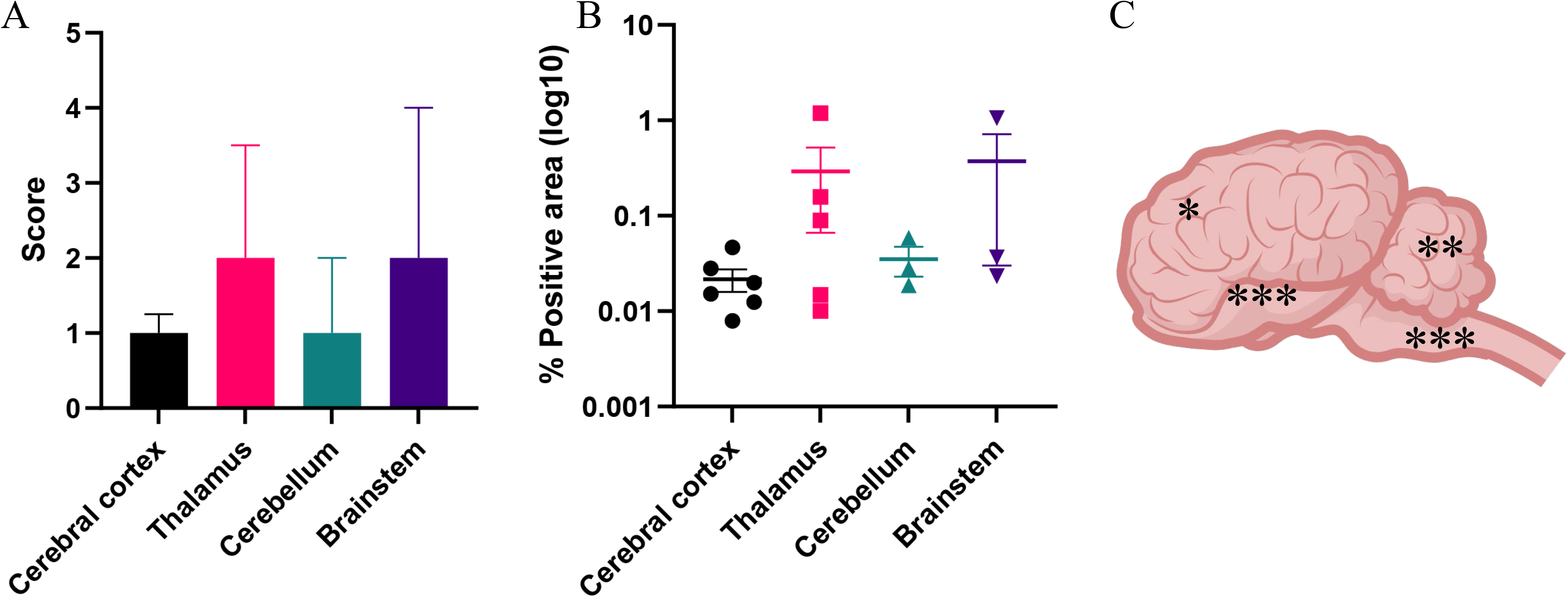
Quantitative analysis of WEEV genomic RNA distribution in the CNS of infected horses. Semiquantitative scores (A) correlate with quantitative pathology analysis (B), with the thalamus and brainstem being the sites with more abundant viral RNA. The mean % positive area was 0.166% (± 0.363%). (C) Graphical view of the equine encephalon. Regions are classified based on the abundance of WEEV RNA in high (***), medium (**) or low (*) based on the quantitative analysis (B).

**Table 3.**
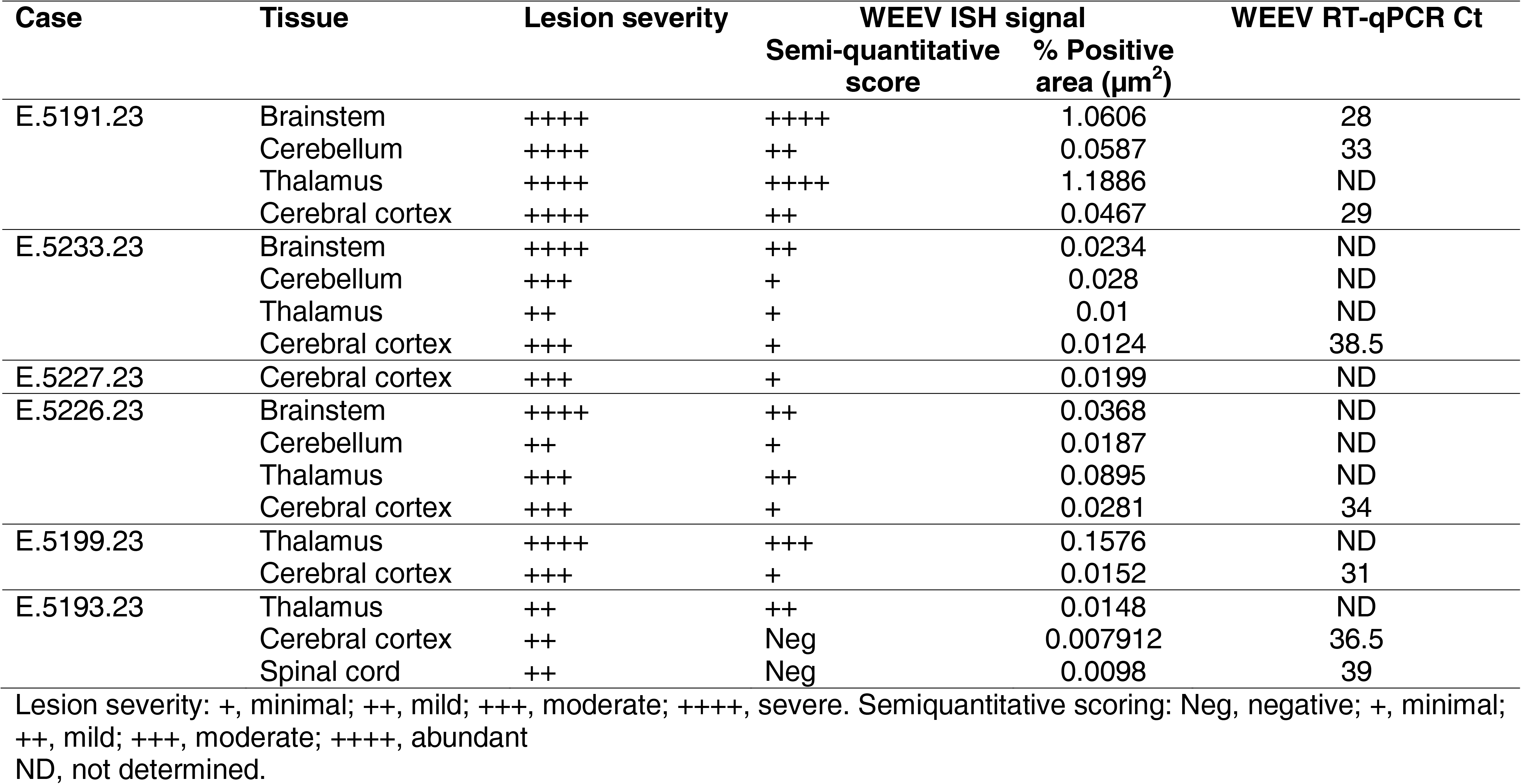
Viral tissue tropism and correlation between histological lesions, ISH signal and Ct values.

## DISCUSSION

Based on the risk assessment published by the Pan American Health Organization (PAHO) as of 14 February 2024, a total of 2,464 outbreaks of WEEV in animals have been officially reported in Argentina (1,445 in 16 provinces), Uruguay (1,018 in 16 departments) and Brazil (single case in one state) [38]. Furthermore, a total of 73 human cases have been confirmed in Argentina (n=69) and Uruguay (n=4), with 7 reported fatalities in Argentina; this number of human cases is significantly higher than that reported in the last epizootic in Argentina between 1982-1983. The incidence rate of the 2023-2024 WEEV outbreak has declined since March 2024 (last case reported on March 7^th,^ 2024). This is probably a result of the implementation of vaccination and establishment of herd immunity, however, underreporting in affected rural areas could be also a factor. Nonetheless, this outbreak has been of higher magnitude compared to previous regional outbreaks and underscores the reemergent potential of this zoonotic disease of public health concern.

Here, we report the virological, serological, pathological and phylogenetic features of cases received at the Equine Virus Unit, INTA during the 2023-2024 WEEV epizootic in the horse population from Argentina. In agreement with recent phylogenetic reports using near full-length genome sequences [39], phylogenetic analysis of partial nsP4 sequences from WEEV isolates obtained from horses derived from three different provinces in Argentina group within the newly proposed lineage C, along with WEEV sequences obtained from Uruguay and Brazil. These findings suggest a common origin for currently circulating WEEV in South America. Interestingly, the lack of closely related isolates between North and South America suggests that several WEEV genotypes are restricted to South America, and such restricted distribution favors the hypothesis that South American WEEV genotypes may be adapted to hosts of restricted geographical distribution [3].

Pathologically, fatal WEEV infections in horses were characterized by often severe meningoencephalitis with an intense neutrophilic component; however, slight variation in the intensity of this inflammatory response was noted depending on the region of the encephalon (Table 3). We demonstrated that inflammatory lesions were most frequent and intense along the internal capsule, thalamus, and brainstem (pons and medulla oblongata) compared to other regions of the brain. Similarly, WEEV RNA abundance was positively associated with the degree of severity of the inflammatory process. Overall, the pathological aspects of WEEV infection in horses, particularly the neutrophilic-type of inflammation, are very similar and indistinguishable to those induced by EEEV, with a similar distribution and abundance of viral RNA in the central nervous system. This underscores the need to include WEEV as a diagnostic differential despite the fact that no outbreaks of disease have been reported in North and South America for approximately 4 decades. Importantly, based on the abundance of WEEV RNA and on its multifocal distribution, our study concludes that collection of fresh specimens from multiple areas of the encephalon, in particular from the brainstem and thalamic region, represents the best sampling approach for maximizing diagnostic success via RT-qPCR. Viral load in the cerebral cortex and cerebellum is often limited and, hence, increasing the number of regions sampled is a beneficial practice to implement. Additionally, lack of detection of WEEV RNA in CSF along with mostly lack of WEEV-specific neutralizing antibodies in this sample type indicates that this is not a reliable sample for antemortem diagnosis of WEEV infection. Among the CSF samples evaluated in this study, a single sample had a high neutralizing antibody titer (1:160). However, no paired serum sample from this horse was submitted and contamination of CSF with blood is possible and could explain the comparatively high levels of antibodies to other samples analyzed.

Despite the limited number of serum samples gathered from horses comingling with WEEV-infected cases, our preliminary serological data demonstrates that over 73% of these animals had virus neutralizing titers against WEEV. These animals were not previously vaccinated against EEEV and WEEV. While no thorough follow-up information was collected from such cases, this serological data suggests that not all equine cases are fatal and rise of specific neutralizing antibodies is noted in surviving animals. Furthermore, it also suggests that the magnitude of the outbreak is likely higher than what may have been reported based on confirmatory testing. Hence, serological tools were critical as an additional diagnostic methodology to complementarily assess the outbreak’s magnitude. However, considering that since January 2024 vaccination against EEEV and WEEV in horses became mandatory in Argentina, there is an urgent need to implement serological tests (e.g. IgM ELISAs) that are able to differentiate vaccinated from natural infected horses for the identification of cases during the acute phase of the disease.

The factors that led to the reemergence of WEEV and current epidemic in South America remain to be determined, but such event is likely multifactorial. There is evidence supporting continuous WEEV circulation within its enzootic cycle in the region [13, 25] and, therefore, factors such as strain variation, ecological and climate change, anthropogenic factors, changes in vector and reservoir dynamics within an immunologically naïve horse population and other factors likely played a significant role.

In conclusion, this is the first description of the virological, serological and pathological features of the 2023-2024 WEEV outbreak in horses from Argentina. Despite its limited occurrence in recent years, WEEV should be considered a differential diagnosis in horses suffering from neurologic disease. This outbreak critically highlights the need for further studies to address ecological drivers determining WEEV re-emergence as a disease of animal and public health concern.

## ACKNOWLEDGEMENTS

We would like to kindly acknowledge Florencia Trubbo, from Garrahan Hospital (Buenos Aires, Argentina) for her excellent technical assistance with tissue processing and slide preparation, the Histology and Immunohistochemistry laboratory at the Louisiana Animal Disease Diagnostic Laboratory (LSU Diagnostics) for their assistance performing RNAscope^®^ ISH and all private veterinarians for collection and shipping of samples, specially Dr. Enrique Durrieu, for collection of paired serum of those convalescent horses during a period of three months.

